# GenVS-TBDB: A Target-Aware AI-Generated and Virtual-Screened Small-Molecule Library for Tuberculosis Drug Discovery

**DOI:** 10.1101/2025.04.28.650819

**Authors:** Xiaoying Lv, Han Guo, JingLan Wang, Qiping Hu, Mengyun Zhang, Heng Wang, Genhui Xiao, Xueying Zheng, Xi Lu, Zhuo Tang, Kai Yu, Gengwei Xu, Shawn Chen, Rumin Zhang, Jinjiang Guo

## Abstract

Tuberculosis (TB) remains a leading global health threat, with over 10 million new cases and 1.25 million deaths reported in 2023. Current TB therapies rely on a limited drug repertoire and prolonged treatment courses that have changed little in four decades. Here, we present GenVS-TBDB, a target-aware AI-generated and virtual-screened small-molecule database, which expands the chemical space against Mycobacterium tuberculosis (M.tb) essential proteins. We first identified 460 probable small-molecule binding pockets across 377 essential M.tb proteins by integrating multiple sources. Then, by leveraging the target-aware molecule generative model, over 1.2 million novel small molecules tailored to these pockets were produced. The key physicochemical properties were computed for all compounds to ensure medicinal chemistry tractability. These compounds were also evaluated using molecular docking and anti-TB specific graph neural network model, yielding binding propensity ranking and whole-cell activity prioritization for each target. To substantiate the integrated AI-driven workflow for anti-TB discovery, 30 compounds were obtained, including 22 AI-designed molecules synthesized de novo and 8 commercially available analogs. In validation, 2 synthesized compounds demonstrated significant thermal stabilization of FtsZ, confirming target engagement. 6 compounds exhibited cellular inhibition below 50 µM, the most potent at 12 µM. Furthermore, GDI-11785 showed binding to the cell wall biosynthesis pathway with 35 µM cellular activity, establishing a promising starting point for tuberculosis drug discovery. Our findings contribute a significant library of bioactive molecules which may hasten the preliminary phase of tuberculosis drug discovery. The GenVS-TBDB small library is publicly accessible and can be downloaded freely at https://datascience.ghddi.org/database/view.

## Introduction

Tuberculosis, caused by Mycobacterium tuberculosis (M.tb), is one of the top infectious causes of death worldwide. Despite decades of control efforts, TB caused an estimated 10.8 million new cases and 1.25 million deaths in 2023^1^. The rise of multi-drug-resistant TB (MDR-TB) further exacerbates the crisis, with ∼450,000 new rifampicin-resistant cases in 2021. Standard treatment for drug-sensitive TB requires a 6-month regimen of multiple antibiotics that has remained essentially unchanged for 40 years^2^. This lengthy therapy and its side effects lead to adherence challenges and treatment failure. Only a few new TB drugs have emerged in recent years – for example, bedaquiline (approved in 2012) was the first new TB drug in over four decades^3^ – and these are mostly reserved for resistant cases^4–6^. Clearly, there is an urgent need to discover novel TB drug candidates that can shorten therapy and overcome resistance.

One promising strategy is structure-based drug design (SBDD), which leverages the 3D structures of essential microbial proteins to guide inhibitor discovery. M.tb has approximately 600 essential genes (roughly 15% of its genome) required for in vitro growth^7^, representing a wealth of potential drug targets across various pathways. With advances in structural biology – including X-ray crystallography and AlphaFold predictive model^8^ – we now have structural data or high-quality models for many of these essential proteins. Structure-based virtual screening can identify small molecules that fit into a target’s binding pocket^9^, enabling rational targeting of enzymes and proteins vital to M.tb survival. However, traditional virtual screening of large chemical libraries is limited by library diversity and size. Here, generative artificial intelligence (AI) can expand the chemical space: generative models can design novel compounds optimized for a given protein pocket, and predictive models can forecast biological activity^10^.

Recent breakthroughs in AI-driven drug discovery have demonstrated the ability to rapidly generate and optimize novel compounds^11–14^. For example, deep generative models have produced potent kinase inhibitors in a matter of weeks^10^. TamGen (i.e., Target-aware molecule generation) is a state-of-the-art target-aware molecule generator that employs a GPT-like chemical language model conditioned on protein pocket context^15,16^. By integrating protein structural information into the generation process, TamGen can propose compounds specifically tailored to a target binding site, improving the likelihood of binding and physicochemical properties. In parallel, AI predictive models like Ligandformer^17^ (a graph neural network originally developed for compound property prediction) enable in silico phenotypic screening, i.e. predicting which compounds are likely to exhibit anti-TB activity. Such models can learn from datasets of active and inactive compounds to prioritize new molecules with desirable bioactivity profiles^18^.

In this work, we present GenVS-TBDB, a target-aware AI-generated and virtual-screened small-molecule database, to accelerate TB drug discovery. We focus on essential M.tb proteins as targets, identifying binding pockets and generating novel small molecules that could potentially bind to these targets. The compounds are assessed for binding affinity, drug-likeness and bioactivity potential (Figure 1). Then, we dual-rank these molecules based on their docking scores and AI-predicted anti-TB cellular activity scores. A total of 22 compounds were synthesized (7 predicted with target activity, 13 predicted with cellular activity, and 2 negative controls) and 8 commercial analogs were purchased, with eight manifesting either target engagement or cellular activity or both, thereby underscoring the robust predictive capability of our artificial intelligence framework. Importantly, we share the GenVS-TBDB, including compounds’ virtual screening analysis data, openly with the scientific community, providing an open resource to jump-start experimental testing and lead optimization. This approach exemplifies how generative AI and computational tools can greatly expand the hit discovery funnel against a high-priority infectious disease.

**Figure 1:**
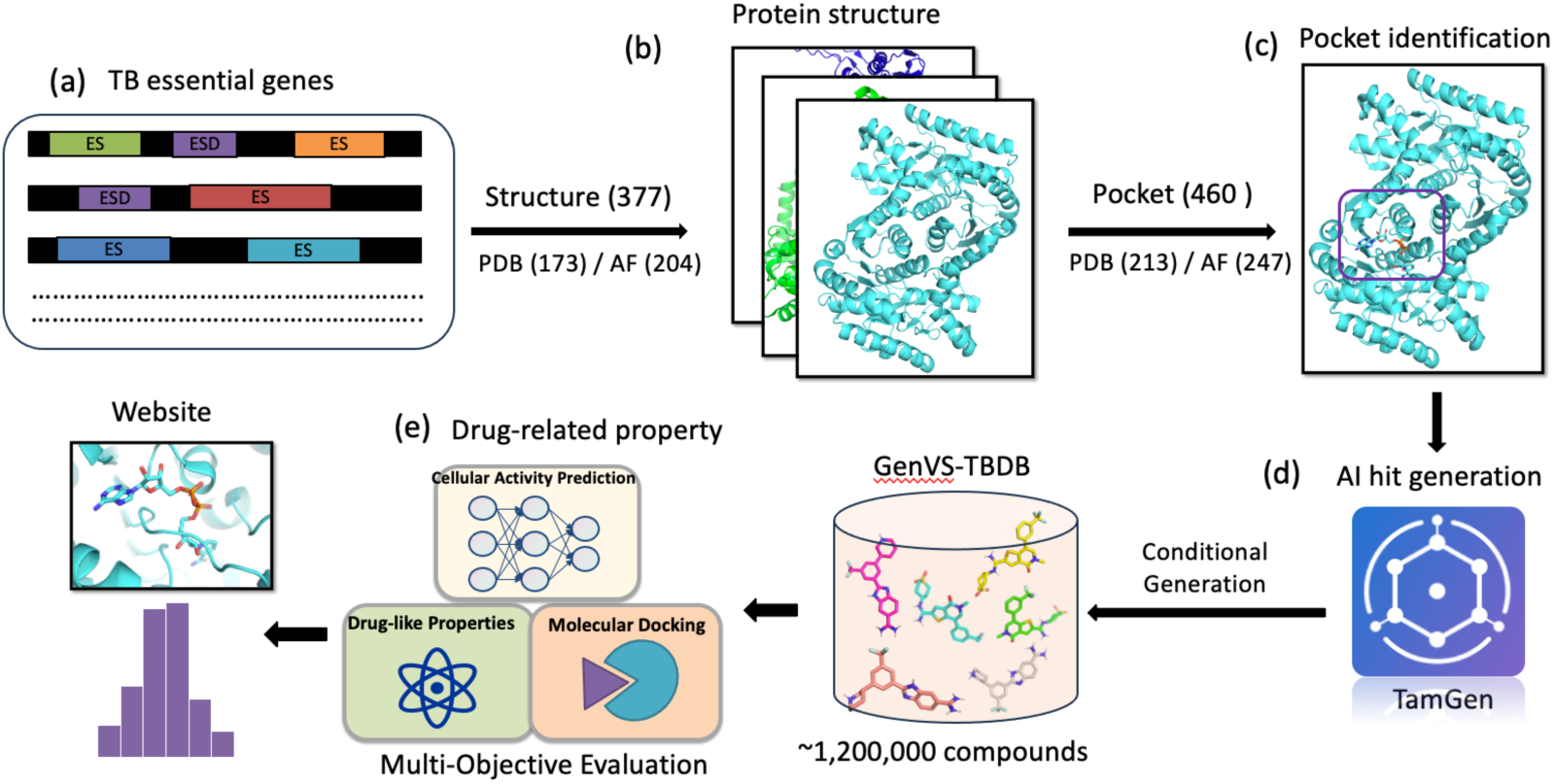
Workflow for constructing GenVS-TBDB. Key steps: (a) selection of an essential M.tb protein target; (b) retrieval or modeling of its 3D structure; (c) identification of binding pockets via co-crystal ligands, homology alignment, and pocket prediction algorithms (Fpocket); (d) compilation of all likely pockets to be used for compound generation; (e) Multi-objective evaluation with molecular docking, AI-predicted cellular activity and drug-like properties for all compounds in GenVS-TBDB.

## Methods

### Target Selection

We selected a panel of M.tb protein targets based on genetic essentiality and druggability. Essential genes were defined according to genome-wide transposon mutagenesis studies^7^, which have identified hundreds of loci indispensable for M.tb growth in vitro. From a comprehensive essential-gene list^7^, we downloaded 1,157 protein crystal structures of these targets from the Protein Data Bank (PDB) database^19–21^. In cases where a target had multiple crystal structures, we selected 173 crystal structures considering their resolution, protein length and presence of bound ligands to ensure reliable pocket identification. For essential targets lacking crystal structures, we harnessed 204 predicted models from the AlphaFold Protein Structure Database, which provides high-confidence structures for most M.tb proteins and allows us to include many otherwise intractable targets^8^. This target selection process yielded a diverse set of proteins, each critical to M.tb survival, forming the basis for downstream pocket identification and ligand generation.

### Binding Pocket Identification

For each chosen target protein, we identified 460 candidate small-molecule binding pockets using a combination of structure-based analyses from highest to lowest confidence level as shown in Table S1:

Known ligand sites: If a target’s crystal structure contained a co-crystallized ligand (such as inhibitor, substrate or analog), we assumed that vicinity as a binding site^22–24^. The bound ligand’s coordinates defined the pocket location and shape. For this category of pocket, a total of 161 binding sites were detected.

Homology-based pockets: For targets without known ligands, we searched for homologous proteins (e.g. in other bacteria H37Ra, M.smeg, E.coli) with ligand-bound structures^25–28^. By aligning the target’s structure with its homolog, we inferred 110 corresponding ligand-binding pockets. Corresponding crystal target proteins were re-constructed using homologous proteins as template to ensure the suitable side chain position in binding pocket.

Binding Pockets Deduced from Literature Review: Our investigation extended to a review of pertinent literature focusing on related crystal structures, with the goal of identifying potential binding sites in the absence of co-crystallized proteins with homology^29–31^. As a result of this comprehensive literary analysis, we pinpointed 20 binding pockets specific to this category.

Computational predictions: For targets devoid of the aforementioned information, we utilized the Fpocket, an open-source pocket detection tool, which is adept at discovering surface cavities that possess the requisite dimensions and shape conducive to ligand binding^9^. Typically, we select one or two pockets per target based on their druggability scores. Employing this technique, we successfully identified 169 pockets.

These targets without obvious cavity in surface were excluded in our protein library. All these sources of information were integrated to build a comprehensive pocket map for each protein (Table S1). In many cases, multiple pockets were identified per protein (e.g. orthosteric active site and secondary/allosteric sites). We catalogued 460 distinct pockets across all targets. Figure 1 illustrates this workflow of binding site identification. Each pocket was annotated with its originating evidence (ligand-bound, homologous, literatures or predicted). This pocket compendium guided our molecular generation, with each pocket considered a separate design scenario for creating tailored compounds.

### Molecular Generation with TamGen

We applied the TamGen algorithm^16^ to design novel small molecules for each identified pocket. TamGen is a deep learning-based generative model that extends a transformer-based chemical language model with protein context encoding. In brief, TamGen’s architecture includes a protein encoder that processes the target pocket information (e.g. sequence of pocket residues or 3D environment features, a VAE-based contextual encoder for compound encoding and seed compound-based refinement and a compound decoder that outputs novel molecular structures in string format (such as SMILES notation). By conditioning the generation on pocket features via a cross-attention mechanism, TamGen produces compounds predicted to fit and interact favorably with the given binding site. This approach effectively biases the chemical output toward each specific target site, unlike generic de novo design.

For pockets with available co-crystalized ligands, we generated novel compounds by conditioning on those ligands. In parallel, for all 460 pockets we randomly sampled 10 compounds from a library consisting of ∼50,000 diverse and druglike compounds and generated compounds based on those seeding compounds. This procedure yielded over 1.2 million unique small-molecule structures overall, with roughly thousands of compounds per pocket on average. Notably, TamGen is designed to optimize multiple objectives simultaneously; beyond just pocket shape complementarity, it considers drug-likeness metrics during generation. The model inherently tends to avoid overly large or undruggable structures, aiming for compounds in a medicinal chemistry-friendly range of size and complexity. The chemical space spanned by these generated molecules is enormous, covering a wide variety of scaffolds, functional group combinations, and three-dimensional geometries not found in existing libraries. By generating this AI-driven library, we circumvented the limitations of predefined screening collections and created bespoke molecules for M.tb targets.

### Molecular Drug-like Properties

All generated compounds were characterized by key physicochemical descriptors relevant to drug-likeness and synthesizability. We calculated properties including partition coefficient (clog P), molecular weight (MW), topological polar surface area (TPSA), hydrogen bond donors/accepters and number of rotatable bonds. These descriptors can set a guideline for oral drug-likeness property (MW < 500, logP < 5, TPSA ≤150)^32^. We evaluated these compounds conformed to established another drug-like parameters, as defined by the Quantitative Estimate of Drug-likeness (QED) criteria^33^. Additionally, we computed the synthetic accessibility (SA) score for each compound with RDKit to gauge how easily the molecule might be synthesized^34^. Where necessary, we applied an SA cutoff to eliminate highly complex structures unlikely to be synthetically tractable. We also quantified per-target molecular diversity using pairwise Tanimoto similarities of Morgan fingerprints (ECFP4). To facilitate visual interpretation of the synthesized compound library, a comprehensive comparative analysis was performed between the compounds generated based on TB targets and those for the CrossDocked2020 test set containing 100 protein pockets, employing the previously described multidimensional evaluation metrics.

Another crucial descriptor was the alert of PAINS (Pan-Assay Interference Compounds) and other frequent hitters. We screened all structures against known PAINS substructures^35^ – chemically reactive or promiscuous motifs that often give false positives in assays. Any compound containing a PAINS alert (such as catechols, aggregating dyes, etc.) was flagged in the database to avoid pursuing problematic chemotypes. By applying these property-based descriptors, we ensured that the final set of proposed molecules not only have high predicted efficacy but also possess favorable drug-like profiles and fewer liabilities. All cheminformatics analyses were performed with standard libraries (e.g. RDKit for property calculation).

### Molecular Docking to Assess Bioactivity Potential

To evaluate the binding potential of the generated molecules, we performed molecular docking for each compound into its designated target pocket. We used AutoDock Vina 1.2.5, a widely used open-source docking program known for its speed and accuracy^36^. Vina 1.2.5 offers enhancements including a reoptimized scoring function and a convenient batch mode for high-throughput docking – critical features for handling our large library. Prior to docking, protein structures were prepared by adding missing hydrogen atoms, assigning protonation states appropriate for pH 7, and defining a docking box around the pocket region. Ligand structures were generated by TamGen in SMILES, which we converted to 3D conformations and minimized. Each ligand was docked into the pocket using multiple runs to sample different binding poses. Vina outputs a docking score (in kcal/mol) for the top poses, which is an estimate of the free energy of binding (more negative indicates stronger predicted binding). The positive docking scores of molecules were set to empty, as they are unlikely to bind to their target pockets.

We recorded the best docking score for each compound–target pair. These scores allowed us to rank compounds and compare the relative druggability of different pockets. As an internal check, for some targets with known inhibitors, we docked the known ligand as a reference. In general, compounds with scores better (more negative) than that of a known moderate inhibitor (∼–7 kcal/mol) were empirically considered as interesting hits. After docking the entire set, we compiled a distribution of docking scores for each target pocket (illustrated in Figure 2a) using **R** (version 4.1) ggplot2 function.

**Figure 2:**
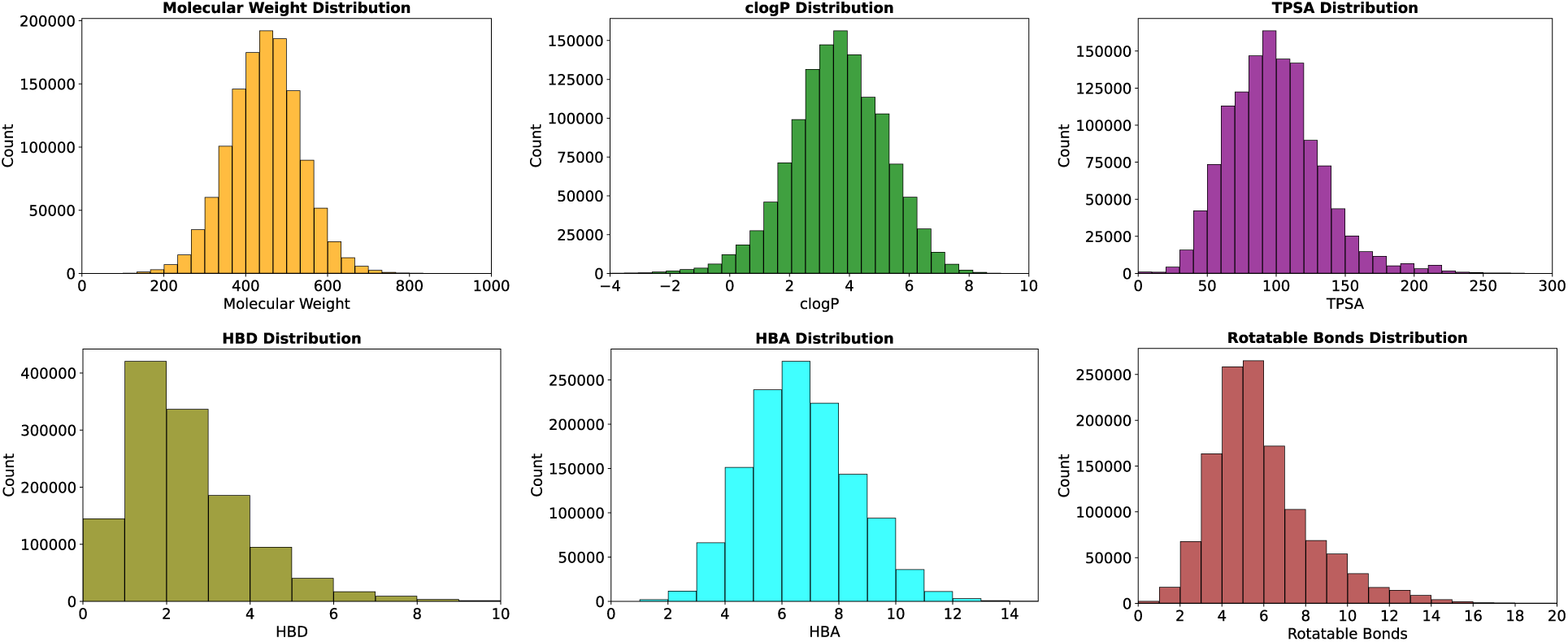
Physicochemical property profiles of generated molecules. Histograms of molecular weight, cLogP, TPSA, hydrogen bond donors/accepters and number of rotatable bonds for the compounds, indicating adherence to drug-like ranges.

### AI-predicted Whole Cellular Activity Model

In parallel with docking-based scoring, we employed an AI-predicted phenotypic activity predictor to further sort the compounds. We used a deep learning model, namely Ligandformer, which is a graph neural network framework tailored to predict compound bioactivity^17^. Specifically, we trained/adapted Ligandformer to predict anti-TB cellular activity, using publicly available and in-house data on known TB-active (Minimum Inhibitory Concentration, i.e., MIC ≤10 µM) and inactive molecules^37^. The training data were collected from multiple sources (Table S2), including internal high-throughput screening, in-house project datasets, and experimental results obtained through collaborations with partners such as GSK. We also incorporated publicly available data where compound activities were reported as AUC values^38^. Based on curve fitting, we determined an AUC threshold corresponding to ∼10 µM MIC and used it as the cutoff to convert AUC values into binary classification labels. The final dataset comprised nearly 5,000 compounds with a balanced positive-to-negative ratio (1:1.08). To ensure robust evaluation, we applied OPTICS clustering^39^ and extracted 500 representative compounds as an independent validation set.

Given a compound’s structure, Ligandformer outputs a probability score estimating the likelihood of achieving an MIC below 10 µM against *M. tuberculosis*. In a three-seed ensemble, the model achieved an average recall of 0.74 and an AUPR of 0.77 at a 0.5 activity threshold on a hold-out test set. On the independent validation set, it achieved an 83% hit rate within the top 100 ranked molecules and maintained ≥70% through the top 200, demonstrating its effectiveness in rapidly surfacing true actives for downstream experimental testing (Figure S1).

### Thermal shift assay (TSA) against FtsZ target

TSA was used to detect protein-compound interactions and performed in a 384-well microliter plate with the Applied Biosystems QuantStudio 7 Real-Time PCR system (Thermo Fisher Scientific, Waltham, MA). The reactions contained 200 μM GDP (Thermo Fisher Scientific, Waltham, MA) and 10X SYPRO Orange in 10 µL MES buffer (50 mM MES, pH6.5, 50 mM potassium acetate, 5 mM magnesium acetate) with three different FtsZ protein concentration 25 μM, 50 μM and 100 μM. Each compound was incubated with FtsZ protein and GDP at room temperature for 2h or 16h, before adding 10X SYPRO Orange. Melting curves were obtained by gradual increase of temperature from 25 °C to 99 °C (0.05 °C increase per second) with fluorescence intensities read at each degree of temperature using optical filters set for carboxyrhodamine (ROX) dye. Melting temperatures (Tm values) were calculated using the Protein Thermal Shift software (Thermo Fisher Scientific, Waltham, MA).

### ATP consumption assay and IC_50_ determination against PheRS target

The assay was performed as previously described^40^. Assays were performed in 384-well white flat-bottom microplates (Corning, #3570). Test compounds were dispensed using the Echo 520 liquid handler across an 8-point concentration series. Wells containing DMSO served as negative controls, while PF-3845 was included as a positive control. Recombinant M.tb PheRS was diluted to 100 nM in assay buffer (50 mM HEPES, pH 7.4, 100 mM NaCl, 10 mM MgCl₂, 0.5 mM TCEP, 0.1 mg/mL BSA, and 0.01% Brij-35) and 5 μL of the enzyme solution was added to each well containing compound. Plates were incubated at room temperature for 30 minutes to allow compound–enzyme interaction. Reactions were initiated by adding 5 μL of substrate mix containing final concentrations of 1 μM ATP, 20 μM L-phenylalanine, and 0.1 mg/mL tRNA^Phe^ in assay buffer. Plates were incubated at 37°C for 2 hours. Following incubation, plates were cooled to room temperature, and 10 μL of 1:50 diluted Kinase-Glo Max reagent (Promega, #V6071) was added to each well. After a 20-minute incubation, luminescence was measured using an EnVision multimode plate reader (PerkinElmer). Data analysis and curve fitting for IC₅₀ determination were conducted using GraphPad Prism 10.

### Thermal shift assay (TSA) against MDH target

In this assay, compounds or DMSO were first transferred into a microplate using an Echo system. Then, 5 μL of tbMDH solution was added to reach a final protein concentration of 4 μM, followed by thorough mixing. Next, 5 μL of diluted SYPRO Orange dye was added to achieve a final 10× concentration, and the mixture was again mixed well. The plate was sealed with a film and briefly centrifuged vertically for 20 seconds to ensure even distribution. Finally, the prepared plate was placed in an ABI Q7 qPCR instrument, and a melting curve program was performed to monitor protein thermal shift.

### Intra-bacterial ATP production assay against cell wall biosynthesis pathway

Intracellular ATP levels were quantified using a luminescent ATP detection assay as previously described^41^. Mycobacterium tuberculosis H37Ra (ATCC 25177) was cultured in liquid Middlebrook 7H9 medium with 10% oleic acid-albumin-dextrose-catalase (OADC), 0.5% glycerol, and 0.05% tyloxapol to mid-log phase, diluted to the desired inoculum, and dispensed into 96-well white plates containing a 10-point, two-fold serial dilution of test compounds. The plates were incubated at 37 °C with 5% CO₂ for 24 h. Following incubation, BacTiter-Glo™ reagent (Promega), equilibrated to room temperature, was added to each well, mixed gently, and incubated for 10 min at room temperature before measuring the relative luminescence units (RLU) using a microplate reader. In parallel, an identical assay plate was incubated under the same conditions for 5 days, after which the bacterial growth was quantified by measuring optical density at 600 nm (OD₆₀₀) and normalized to the DMSO control.

### Whole Cellular Activity in H37Rv strain

M.tb H37Rv (ATCC27294) strain was cultured in Middlebrook 7H9 Borth (Becton-Dickinson, Franklin Lakes, NJ) with 10% OADC (Becton-Dickinson, Franklin Lakes, NJ), 0.05% tween-80 at 37 °C, 5% CO2 to an optical density at 570 nm (OD_570_) of 0.4-0.7. Compounds were added into 96-well plate (Becton-Dickinson, Franklin Lakes, NJ) by two-fold dilution from 200 μM in 100μL medium. H37Rv cells were diluted to OD_570_ of 0.002, which corresponded to 2 x 10^5^ CFU/mL in this case, and seeded to each compound concentration in 100 μL medium per well. The final highest concentration of compounds was 100 μM and the final concentration of cells was 1 x 10^5^ CFU/mL. The plate was incubated for 7 days at 37 °C and 5% CO_2_. Subsequently mixture of 20μL Alamar Blue (Thermo Fisher Scientific, Waltham, MA) and 12.5μL 20% tween-80 was added per well and incubated the plate for another 24 hours. The fluorescence signals of emission wavelength at 590 nm for each well were recorded, and H37Rv MIC was defined as the lowest concentration where inhibition of fluorescence signal is greater than or equal to 90%.

### Cytotoxicity assay in Vero cell line

Vero cells are cultured in ATCC DMEM medium supplemented with 10% fetal bovine serum (FBS) and Penicillin/Streptomycin. The cells are incubated in a humidified atmosphere of 5% CO*_2_* at 37°C and then diluted to 1×10*^5^* cells/ml. The drugs were diluted in DMSO and serially two-fold diluted in 96-well plates from 100 μM. After incubation at 37°C for 48 hours, cytotoxicity is tested by CellTiter-Glo luminescent cell viability assay (Promega, Beijing, China) and the luminescence signal was recorded by SpectraMax-M5 plate reader (Molecular Devices, San Jose, CA). IC_50_ were determined using the non-linear fit (log(inhibitor) vs. normalized response Variable slope) function in Prism 8.

### Binding likelihood evaluation of kinases from Vero cell line

To elucidate the cytotoxic nature of our compounds, we identified all kinases protein (uniport ID: A4KY21, B9UKH5, P10650, Q14UF0, Q14UF1, Q50GY5, Q9GK79, Q9GMA1, Q9GMA2, W0GDK6, W0GHV6) from the Vero cell proteome by searching the UniProt database with the key terms ‘Chlorocebus aethiops’ (Green monkey)^42^. The probabilistic assessment of ligand-target interactions for the synthesized compounds was computed utilizing Boltz-2, a cutting-edge computational model for biomolecular simulations. This tool is at the forefront of integrating deep learning methodologies to approach the accuracies yielded by established physics-based free energy perturbation (FEP) techniques, initially developed by researchers at the Massachusetts Institute of Technology (MIT)^43^. Notably, Boltz-2 has been reported to offer orders-of-magnitude computational advantage compared with traditional FEP methods. This advancement substantially enhances the practicability of precise in silico screening processes vital for the preliminary phases of drug discovery.

## Results

### GenVS-TBDB Library Drug-Likeness Properties

In this study, a 1.2 million-level small molecule library GenVS-TBDB was generated for about 400 TB essential targets. We calculated the physicochemical profiles of the library to evaluate whether they meet desirable drug-like criteria. Figure 2 depicts the distributions of key molecular properties--molecular weight, cLogP, topological polar surface area (TPSA) hydrogen bond donors/acceptors and rotatable bonds--for all compounds after property calculation. The compounds span a range of sizes: molecular weight ranges from about 250 up to 500+ Da, with a median around 360 Da. This is within a typical small-molecule drug range and suitable for potential oral bioavailability. The clogP (calculated octanol-water partition coefficient) distribution centers around 3–4, with the majority between 1 and 5 log units, indicating moderate lipophilicity. Only a small fraction has clogP > 5, which could signal solubility issues. The topological polar surface area (TPSA) values cluster in the 50–100 Å^^2^ range, consistent with good permeability (generally TPSA <140 Å^^2^ is favorable for oral drugs).

We also compared the drug-like properties of molecules of the GenVS-TBDB with those from the CrossDocked2020 test set^44^, using metrics such as QED, molecular diversity, synthetic accessibility (SA), and Lipinski rule compliance. For instance, the GenVS-TBDB molecules achieved a molecular diversity score 0.866, outperforming the CrossDocked2020 test set (Table 1). The synthetic accessibility (SA) score, reflecting favorable synthetic feasibility, was calculated as 0.782 for our dataset, which is comparable to or exceeds values reported for other reference library. In contrast, the quantitative estimate of drug-likeness (QED) and Lipinski’s rule compliance scores were lower in our dataset (0.497 and 0.607, respectively) compared to the reference dataset (QED = 0.559; Lipinski’s rule compliance = 0.988). These results highlight opportunities for further optimization of drug-like properties while retaining robust synthetic tractability in our library.

**Table 1.** Comparison of molecular properties (QED, diversity, SA, Lipinski compliance) between the GenVS-TBDB and other datasets. The highest score in each column was highlighted by bold.

|  | **QED** | **Diversity** | **SA** | **Lipinski** |
| --- | --- | --- | --- | --- |
| **CrossDock2020** | **0.559** | 0.747 | 0.771 | **0.988** |
| **GenVS-TBDB** | 0.497 | **0.866** | **0.782** | 0.607 |

Notably only 0.26% of the compounds in our library exhibited structural motifs associated with known PAINS substructures (Table S3). In comparison to established libraries containing FDA-approved drugs or clinical-stage candidates, the molecules by TamGen demonstrated a PAINS rate of 0.26% (3298 compounds), which is significantly lower than those of the FDA Library (2.6%) and ReFrame Library (2.1%). This stark contrast underscores the reduced risk of promiscuous binding in GenVS-TBDB, thereby enhancing their reliability and prioritization potential for drug discovery pipelines.

### Docking Score Distribution across Targets

This study evaluated the Vina docking score distributions across 1.2 million small molecules screened against 460 essential protein pockets. The analysis identified distinct patterns in ligand-target interactions, with significant variations in binding affinities observed across different pockets. We summarized the distribution of docking scores (AutoDock Vina scores, in kcal/mol; more negative is better) aggregated over all targets to see overall trends as shown in Figure 3a. The distribution is roughly normal: a majority of compounds achieved moderate scores around –6 to –10 kcal/mol (comparable to micromolar to sub-micromolar affinities). The overall median docking score was approximately –7.5 kcal/mol. More than half of the targets had at least one compound with a docking score better than –10, suggesting that TamGen successfully found high-affinity chemotypes for many essential proteins. This indicates that for most target pockets, TamGen was able to generate some ligands with reasonable predicted affinity.

**Figure 3:**
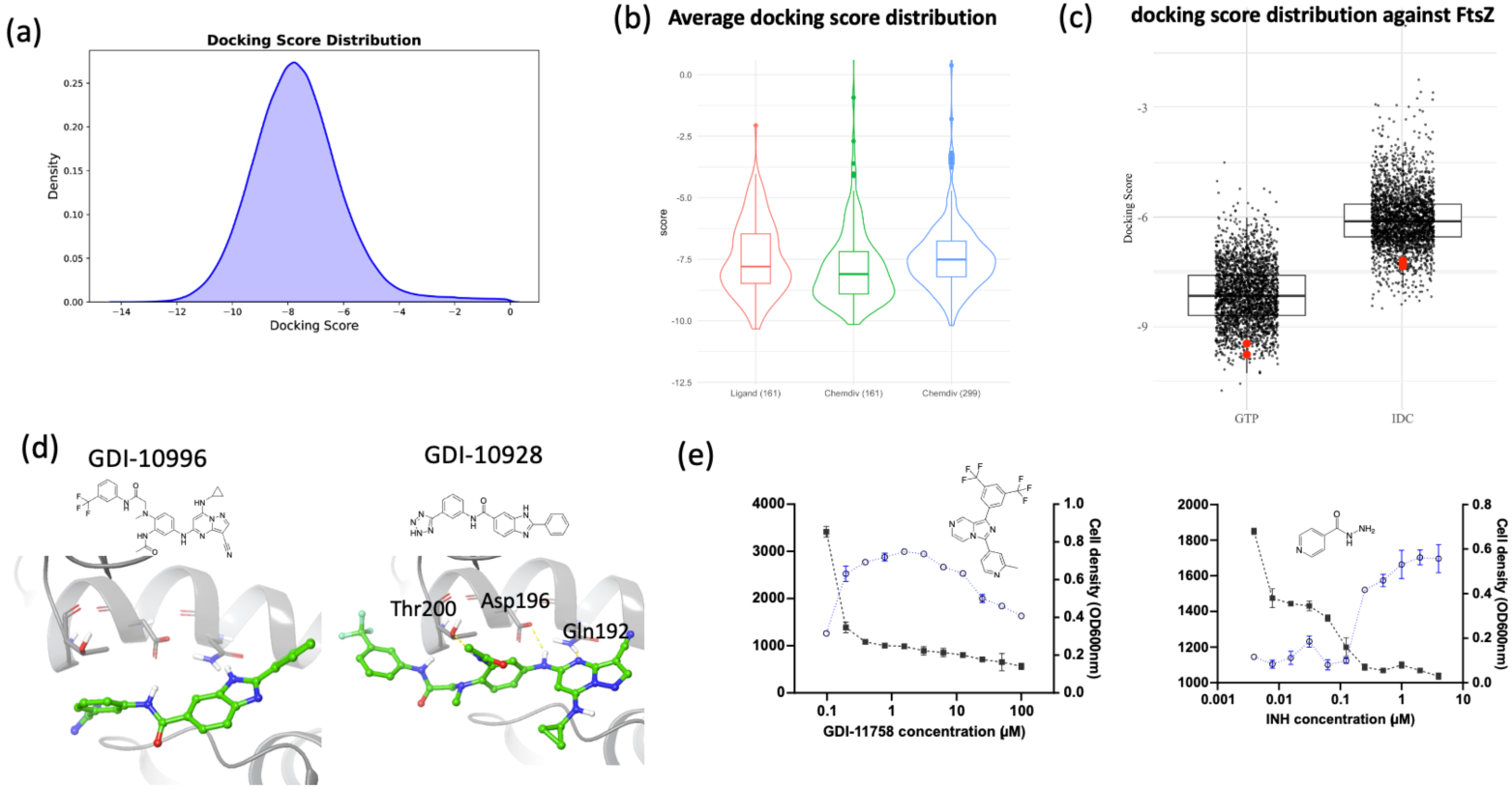
Distribution of docking scores for AI-generated compounds against M.tb targets and related compounds against FtsZ target. (a) The histogram shows the frequency of AutoDock Vina scores across all ∼1.2 million compound–target docking simulations. A lower (more negative) score indicates a better predicted binding affinity. (b) An inset compares score distributions for different target classes (co-crystal proteins vs. apo proteins). Vina average docking scores across three distinct experimental groups. The 161 and 299 in bracket represents these co-crystallized and apo essential proteins. “Ligand” and “Chemdiv” represents the seed of TamGen from co-crystal ligands and chemdiv library, respectively. (c) Distribution of docking scores for related compounds against two FtsZ target pockets. The red dot means the synthesized compound. (d) The binding poses of compound GDI-10996 and GDI-10928 against FtsZ IDC pocket. with h-bond shown by yellow dashed line.

This study also evaluated the Vina average docking scores across three distinct experimental groups: (1) 161 proteins using native co-crystal ligands as seed molecules for TamGen, (2) the same 161 proteins seeded with ten ChemDiv-derived small molecules, and (3) 299 apo proteins similarly seeded with ten ChemDiv compounds. Notably, the second group exhibited the lowest average docking scores (Figure 3b), suggesting enhanced binding affinities compared to the co-crystal ligand group and the apo protein cohort. This disparity may arise from the ChemDiv compounds in the second group adhering more closely to the Rule of Five, compared with the native co-crystal ligands (Group 1), while the apo proteins (Group 3) may accommodate diverse ChemDiv molecules due to their unoccupied, flexible binding sites. These findings underscore the interplay between ligand physicochemical properties and protein conformational states in docking performance, highlighting the need for context-specific compound libraries in structure-based drug discovery. We also note that the distribution of top scores correlates with known target druggability; pockets known to bind potent inhibitors (like the proteasomal enzyme active site) showed tighter score ranges, whereas more challenging or novel pockets had broader, lower-affinity distributions.

At the per-target level, we observed that certain targets yielded consistently better docking hits. The top three representative targets are RmlB (-10.19), Glf (-10.14), EmbC (-10.10) with best average docking score. The broad molecular binding observed in M.tb essential targets—RmlB, Glf, and EmbC—stems from their critical roles in bacterial survival and structural adaptations to diverse substrates. Enzymes such as RmlB (cell wall biosynthesis) exhibit flexible substrate-binding pockets to accommodate nucleotide sugar derivatives, while glycosyltransferases (Glf, EmbC) require tolerance for branched polysaccharide precursors, enabling interactions with varied ligands. Evolutionary pressures under host stress conditions may have further selected for structural plasticity in these targets, combining conserved catalytic cores with peripheral flexibility. This functional and structural adaptability highlights their vulnerability to inhibition but underscores the need for precision in drug design to balance potency and selectivity. These findings position these targets as high-priority nodes for anti-TB therapeutic development.

Conversely, targets Rv1110 (11.38), Rv1990c (31.15), and CarA (32.17) displayed the highest (least favorable) scores, indicating potential challenges in ligand binding or suboptimal pocket geometries. For these three targets with the poorest performance, the limited space within their binding pockets presents a consistent obstacle for accommodating small molecule docking, as illustrated in Figure S2.

These docking results validate the target-aware generation strategy: by designing compounds specifically for each pocket, we achieved good docking fits for many targets. This provides insight into which target classes might be most effectively addressed by our compounds. Importantly, docking alone is an imperfect predictor of true binding affinity; nonetheless, the scoring step was invaluable for cutting down the enormous library to a manageable list of high-scoring candidate molecules for each target.

### Binding affinity validation against three targets

FtsZ, a prokaryotic analog of the eukaryotic tubulin, is indispensable to bacterial cytokinesis, orchestrating cell division by assembling into a Z-ring structure. This fundamental role makes FtsZ a prime candidate for TB drug discovery efforts. Our study, facilitated by the TamGen computational platform, has identified two critical binding domains within the FtsZ protein: the guanosine triphosphate (GTP) binding pocket and the interdomain cleft (IDC). A targeted compound library was generated, comprising 2,684 molecules for the GTP site and 2,757 for the IDC site. Comparative docking analyses, as delineated in Figure 3c, revealed a lower (more negative) average docking score of -8.12 at the GTP site versus -5.99 at the IDC site, indicating a higher affinity of the screened molecules for the GTP pocket. This differential binding efficacy underscores the therapeutic potential of targeting the GTP site of FtsZ in the development of new anti-TB agents.

To validate the binding affinity of FtsZ target, five compounds were picked out mainly based on docking score (cutoff: -7.0) and drug-like properties. The criteria for selection included considerations of molecular stability, metabolic profiles, ease of synthesis, docking scores, among other physicochemical properties. From these five candidates, two compounds, GDI-10928 and GDI-10996, exhibited binding affinity to FtsZ, with thermal shift values comparable to the established control compound GTP. GDI-10928 demonstrated a concentration-dependent thermal shift at 16 hours post-incubation, with increments of 1.84°C, 1.56°C, and 1.23°C at concentrations of 100μM, 50μM, and 25μM, respectively. In contrast, no significant concentration-dependent effects were observed at 2 hours, with thermal shifts recorded as 1.70°C, 1.55°C, and 1.63°C, at the same respective concentrations. Conversely, compound GDI-10996 revealed a clear concentration-dependence both at 2 and 16 hours, as shown in Table 2. At 2 hours, temperature shifts of 0.89°C, 0.67°C, and 0.37°C were observed for concentrations of 100μM, 50μM, and 25μM, respectively, while at 16 hours the shifts were measured at 1.25°C, 0.96°C, and 0.59°C for the same concentration gradient. In our prediction, GDI-10928 can generate three h-bonds with Gln192, Asp196 and Thr200 at IDC pocket, while there is no obvious interaction for GDI-10996 (Figure 3d). These may be the main reason why GDI-10928 compound has higher Tm shift compared with GDI-10996 in our TSA assay.

**Table 2:** The TSA validation against FtsZ target. Two compounds with highest Tm shift were highlighted by bold.

| GDI number | Structure | TSA ΔT<sub>m</sub> (°C), | TSA ΔT<sub>m</sub> (°C), | Vina score |
| --- | --- | --- | --- | --- |
|  |  | 100/50/25 µM 2h | 100/50/25 µM 16h |  |
| **GTP**<br>**(dTmd +)** |  | 4.59/3.60/2.81 | / | / |
| GDI-10927 |  | 0.51/0.22/0 | 0.52/0.22/-0.22 | -7.19(IDC) |
| **GDI-10928** |  | **1.70/1.55/1.63** | **1.84/1.56/1.23** | -7.35(IDC) |
| **GDI-10996** |  | **0.89/0.67/0.37** | **1.25/0.96/0.59** | -7.24(IDC) |
| GDI-11043 |  | -0.37/-0.30/-0.07 | -0.52/-0.37/-0.07 | -9.47(GTP) |
| GDI-11095 |  | -0.15/-0.07/0.07 | -0.22/-0.07/0 | -9.77(GTP) |

The obvious temperature shift and dose response effect, coupled with comparable binding geometries, underscore the potential of GDI-10928 and GDI-10996 as pormising FtsZ inhibitors for the development of novel anti-TB therapeutics. This work provides a foundation for developing novel therapeutics targeting FstZ, although further in vitro and in vivo investigations are warranted to establish the compounds’ efficacy and safety profiles.

Our study also encompassed validation of the binding affinities of two compounds, specifically targeting the essential enzymes phenylalanyl-tRNA synthetase (PheRS) and malate dehydrogenase (MDH). Despite these efforts, minimal inhibitory activity was not observed for PheRS upon evaluation with the ATP consumption assay; compounds GDI-10928 and GDI-10931 did not exhibit notable inhibition. The half-maximal inhibitory concentration (IC_50_) for GDI-10928, as depicted in Supplementary Figure S3, was determined to be 120 μM.

For the MDH target, the Thermal Shift Assay (TSA) was employed to corroborate the binding affinity of the proposed compounds. One compound, GDI-10994, had to be excluded from analysis due to interference from excessive background noise during the assay. GDI-10931 exhibited a negligible thermal shift of 0.3°C at a high concentration of 400 μM, which suggests an absence of significant binding affinity compared with 4.7 degrees shift of the control molecule NADH (Table S4). These results necessitate further investigation to clarify the relevance of the observed effects and to determine the potential of these compounds as inhibitory agents against PheRS and MDH.

### Cellular activity validation in H37Rv strain

Integrating the AI-based anti-TB predictions with the docking results substantially improved our hit prioritization strategy. After obtaining docking scores, we evaluated each compound using the Ligandformer model to predict its anti-TB activity probability. The model outputs a score between 0 and 1 with higher values indicating greater a likelihood that a compound will inhibit M.tb growth, based on patterns learned from known actives. We applied a conservative threshold, tuned on a validation set of compounds with experimentally determined activities, to classify compounds as "predicted active". Rather than reflecting true biological hit rates, this classification was intended to support prioritization and enrichment of candidate molecules. Using this threshold (predicted probability > 0.5 for a cellular MIC ≤ 10 µM), a substantial fraction of the generated compounds were ranked as high-priority candidates by the model (Figure S4).

To evaluate the Ligandformer model’s predictive performance, a set of 15 compounds complemented by eight commercially obtained analogs was curated, principally based on their predicted cellular activity. The selection considered such as chemical diversity, predicted metabolic profiles, ease of synthesis, and other pertinent physicochemical attributes discussed in the previous section. We subsequently measured the minimum inhibitory concentrations (MICs) of these compounds against the H37Rv strain of M.tb (Figure 4).

**Figure 4.**
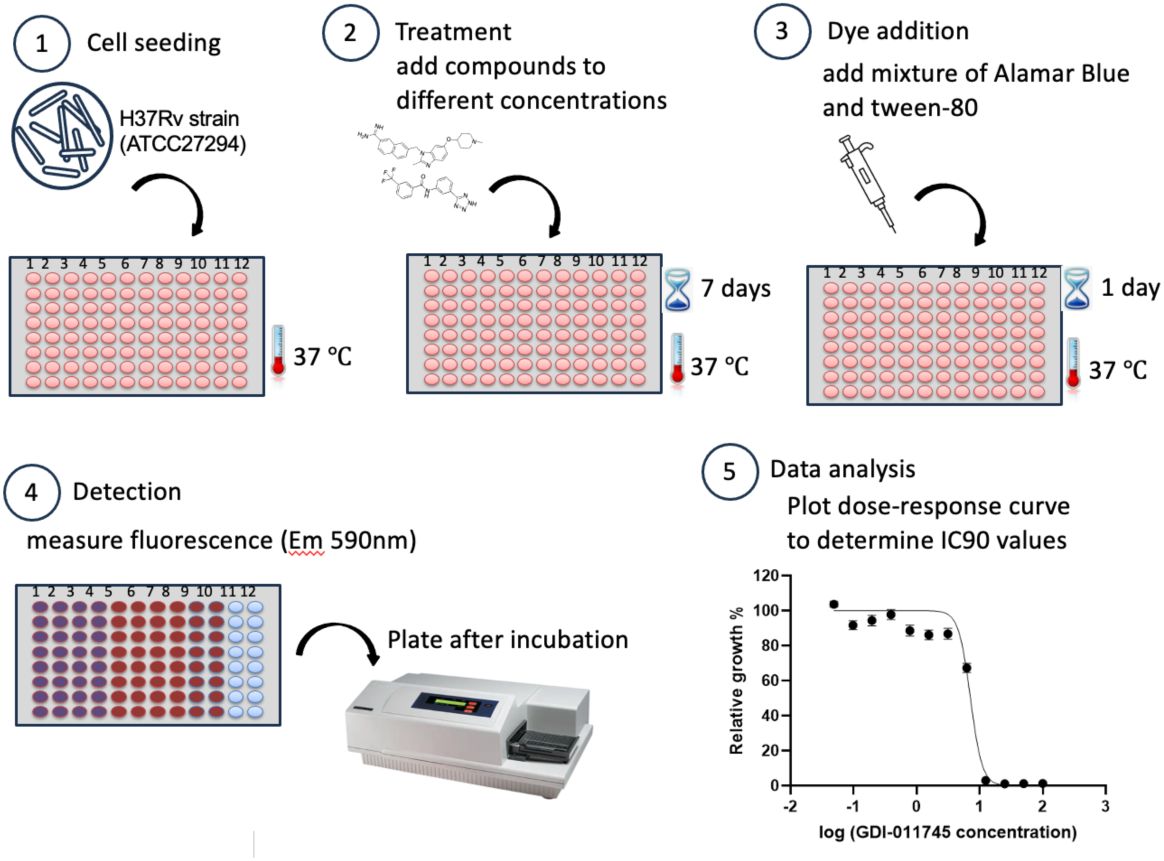
The MIC validation workflow in H37Rv strain. M. tb H37Rv (ATCC27294) strain was cultured at tempature 37°C. Compounds were added into 96-well plate to incubate 7 days at different concentrations. Dye was added per well and incubated the plate for 24 h. The fluorescence signals were recored and H37Rv MIC was defined from relative growth curve.

For the eight purchased analogs, all kinase inhibitors, one analog with the CAS ID 1242156-23-5 revealed the best MIC of 25 µM, although the initial generated molecule GDI-11786 shows no cellular inhibition in H37Rv strain (Table 3). Among these synthesized compounds, five, highlighted by yellow in Table 3, exhibited cellular activity with inhibitory concentrations below 50 µM, with compound GDI-11745 demonstrating potent activity at a concentration of 12 µM against the H37Rv strain as shown in Figure 4. This compound had a predicted probability of displaying cellular activity below 10 µM of 1.0, highlighting the predictive strength of our deep learning algorithm. Compounds GDI-11933, GDI-11785, GDI-11747, and GDI-12005 exhibited MIC of 22 µM, 35 µM, 36 µM, and 45 µM, respectively. Notably, GDI-11785 also demonstrated target engagement with MmpL3 in the ATP-production assay (Figure 3e), indicating that it may serve as a promising starting point for tuberculosis drug discovery. In contrast, compounds GDI-11784 and GDI-11825 were synthesized as negative controls based on predicted cellular-activity probabilities of 0.46 and 0.51, respectively, and both displayed MIC values exceeding 100 µM, highlighted by green in Table 3. This outcome further supports the predictive accuracy and robustness of the computational model.

**Table 3:**
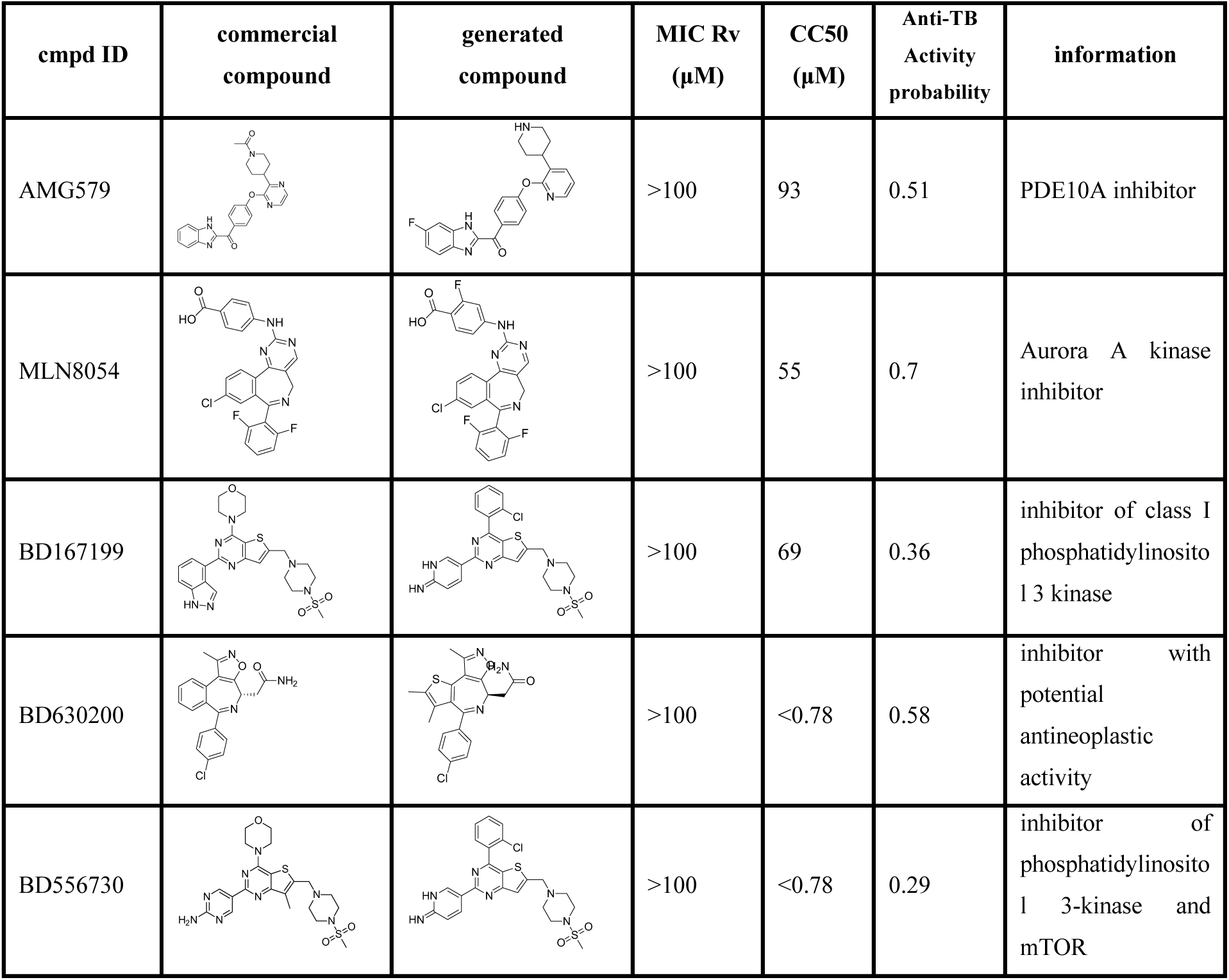
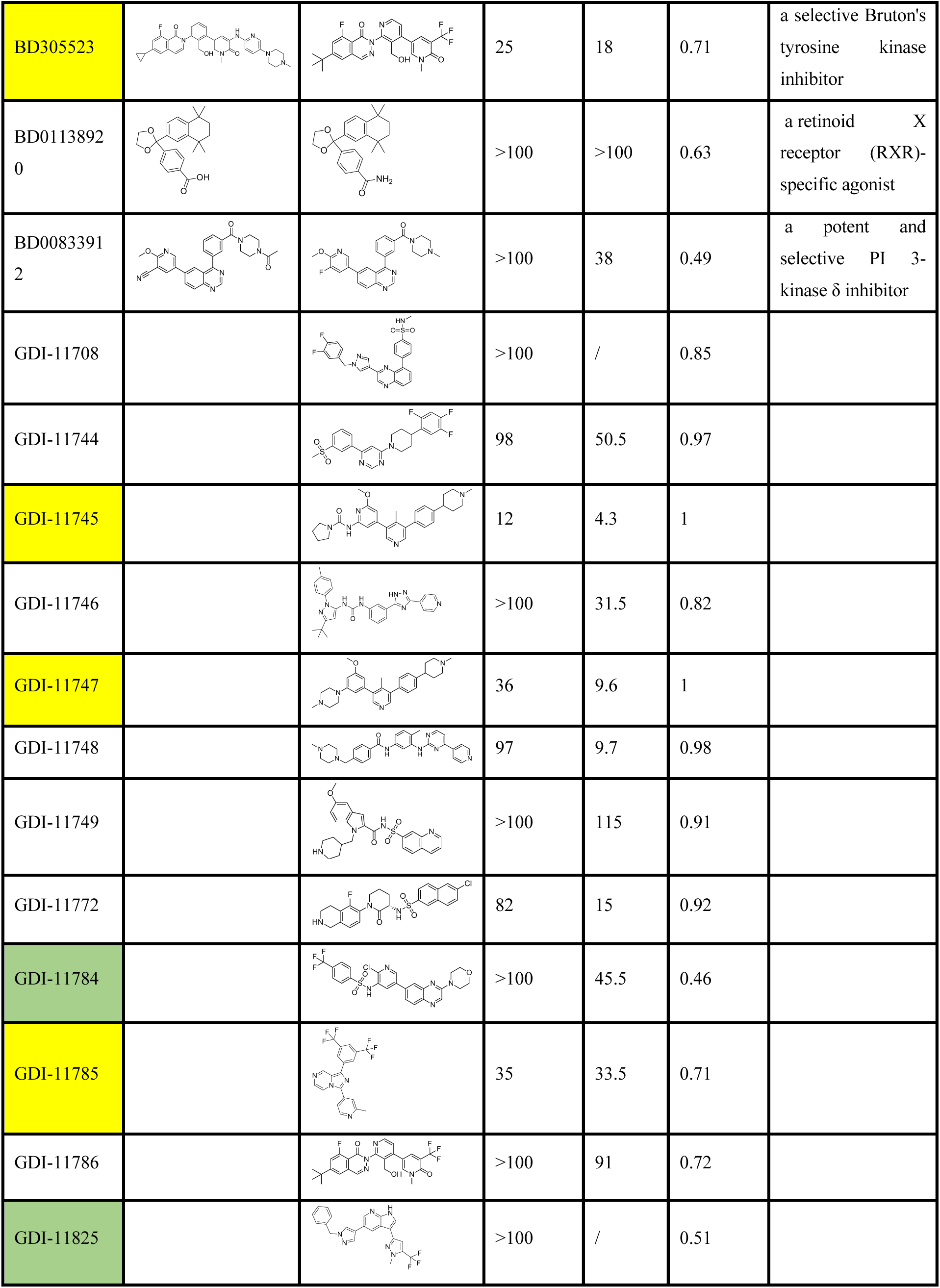
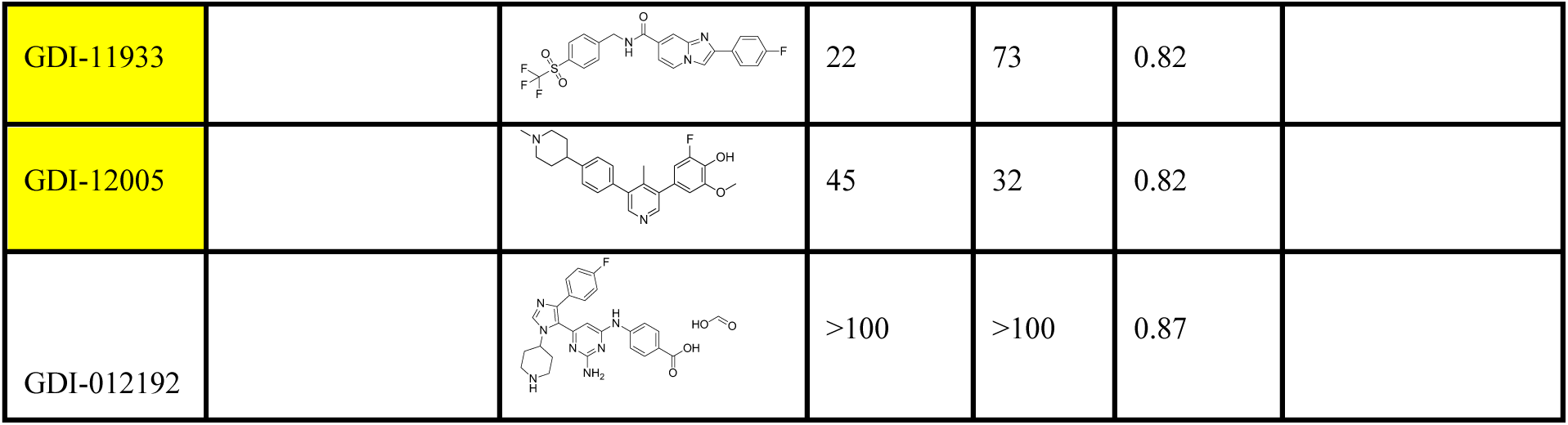
The anti-TB cellular activity validation at H37Rv strain. Six active compounds with cellular activity and two negative controls were highlighted by yellow and green, respectively.

To further probe the potential mechanism of action, we evaluated the binding of the GDI-11785 compound using an intra-bacterial ATP production assay. MmpL3 is an essential membrane transporter responsible for exporting mycolic acid precursors required for mycobacterial cell-wall biosynthesis. As shown in Figure 3e, GDI-11785 increased intracellular ATP levels at concentrations ranging from 0.1 to 1 μM, exhibiting a pattern comparable to that of the anti-TB drug isoniazid (INH), which elicits a similar ATP-enhancing effect within the same concentration range. These findings indicate that GDI-11785 interacts with the cell wall biosynthesis pathway, including MmpL3 target.

Despite these encouraging mechanistic signals, most of these selected compounds, inclusive of six identified as cellularly active, displayed cytotoxic effects on the Vero cell line, as delineated in Table 3. Notably, compound GDI-11745 exhibited a CC_50_ (50% cytotoxic concentration) for the Vero cell line at 4.3 µM, which is lower than its MIC value of 12 µM against the H37Rv M. tb strain. In fact, our analysis unexpectedly revealed that several generated molecules appeared across multiple targets and exhibited favorable predicted binding scores (Table S5), indicating potential multi-target engagement, which may in turn account for their observed cytotoxicity.

Furthermore, to elucidate the cytotoxic nature of our compounds, the binding likelihood were calculated using boltz-2 against the target with highest vina docking score and 11 kinases from the Vero cell line. The results showed that the binding likelihood to the best target was reduced for the synthesized compounds relative to those against the kinases, which suggest these compounds are more likely to interact with the kinases to cause the obvious cytotoxicity in Vero cell line (Figure S5).

### Open-access of the GenVS-TBDB

The GenVS-TBDB (accessible at https://datascience.ghddi.org/database/view) provides a comprehensive repository of M.tb essential proteins, their structural data, and associated small-molecule interactions. Users can search for targets using three primary identifiers: (1) locus name (e.g., Rv3280), (2) gene name (e.g., accD5), or (3) structural ID (e.g., PDB ID 2BZR). Upon querying, the platform dynamically displays interactive visualizations, including 3D protein structures and center position coordinates, and predicted cellular activity profiles of associated compounds. For example, searching for the structural ID "2BZR" directs users to a dedicated page (https://datascience.ghddi.org/database/detail?db=MTB&pid=2BZR) showcasing the crystallographic structure of AccD5 (Rv3280), annotated pockets with spatial coordinates, and a list of docked molecules ranked by Vina scores.

The database supports bulk downloads of all target metadata (e.g., locus tags, gene names, structural IDs, pocket counts) in CSV formats. Additionally, individual target pages enable users to download detailed molecular datasets, including compound smiles physicochemical properties (e.g., molecular weight, LogP, HBA, HBD), and docking scores. This feature facilitates downstream analyses such as structure-activity relationship (SAR) studies or virtual screening workflows.

The integration of structural biology data with cheminformatics tools positions this resource as a critical platform for TB drug discovery, enabling researchers to prioritize targets, optimize lead compounds, and explore novel binding mechanisms.

## Discussion

After downloading data from our service, the user can dual rank the compounds according to their docking scores and predict cellular activity probabilities as prioritization guide. Despite multiple rigorous TSA assay and potency experiment validations, we still treated the AI output as a guide rather than a strict rule – some compounds we set aside as “predicted inactive” might still be worth testing if they have exceptional docking and properties. In summary, the AI-predicted activity and docking prioritized a refined set of high-potential hits: molecules that not only bind well to their intended M.tb target but also carry a computational imprimatur of likely whole-cell efficacy. These prioritized compounds form the basis of our top candidate selection in the subsequent analysis or experimental testing.

Despite the powerful integration of methods, some limitations of our study must be acknowledged. Only one structure for most essential targets was selected for molecular generation. It’s important to note that variations in side-chain positions within the same pocket, as well as different protein conformations, were not taken into account. Meanwhile, docking scores were used for relative ranking rather than absolute affinity estimation.

Our GenVS-TBDB provides fresh chemical matter against both the classic targets and entirely new ones. Notably, many of our top compounds have novel scaffolds unrelated to existing TB medications. The study demonstrates a novel convergence of structure-based methods and generative AI design to address the urgent need for new tuberculosis therapeutics. By targeting a wide array of essential M.tb proteins, we aim to expand the therapeutic arsenal beyond the current limited set of TB drugs. The results yielded multiple potential drug candidates worthy of experimental follow-up.

## Conclusion

Targeting 460 druggable binding pockets on 377 essential TB proteins, we curated a vast chemical library of novel small molecules for TB drug discovery, namely GenVS-TBDB, enabled by an innovative generative AI and virtual screening workflow. The molecular generation model, TamGen, produced over 1.2 million small molecules, which were systematically assessed for binding affinity using docking evaluations and further enriched for anti-TB cellular activity with the AI predictor Ligandformer. For the 22 synthesized compounds and 8 commercial analogs, eight top candidates were experimentally supported with either good target binding affinity or high cellular potency in experiments or both, especially for GDI-11785 with both target engagement against the cell wall biosynthesis pathway and whole-cellular activity at 35 µM and GDI-11745 with more potent activity at 12 µM. In an unprecedented move, we have made this entire library freely available in a public website to foster global collaborative research and expedite TB drug R&D, believing that shared data can spur discoveries by multiple groups simultaneously.

Our work demonstrates the power of generative AI and multi-objective virtual screening in uncovering efficacious anti-TB agents and offers a framework that can extend to other infectious diseases. The next phase involves experimental validations, while the larger vision is to leverage this AI-driven method to address other formidable pathogens, thereby progressing toward ending the TB epidemic through open science and computational advances.

