## supporting figures and tables for "GenVS-TBDB: A Target-Aware AI-Generated and Virtual-Screened Small-Molecule Library for Tuberculosis Drug Discovery"

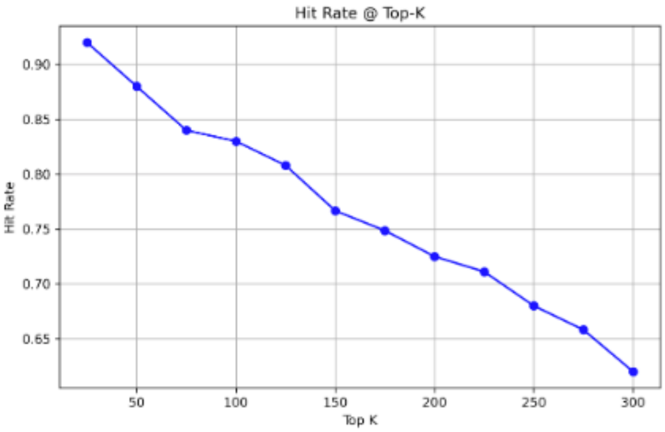


**Figure S1**. The performance of Ligandformer model in predicting anti-TB cellular activity. Hits enrichment on independent validation set, measured with precision@Top-K (random-screening baseline hit rate 23.6%).

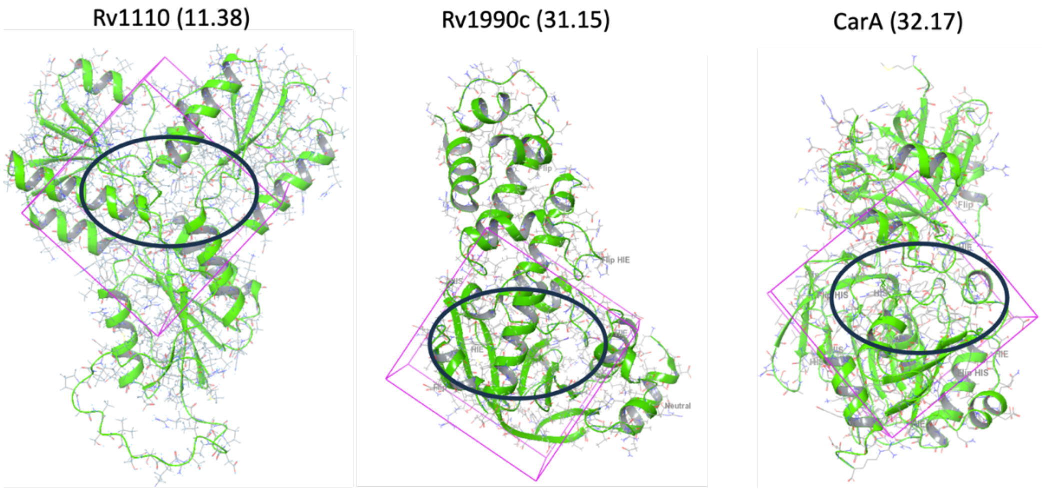


**Figure S2**. The pocket position for three worst targets in average docking score. The pocket position was highlighted by pink and black

GDI-10928 GDI-10931

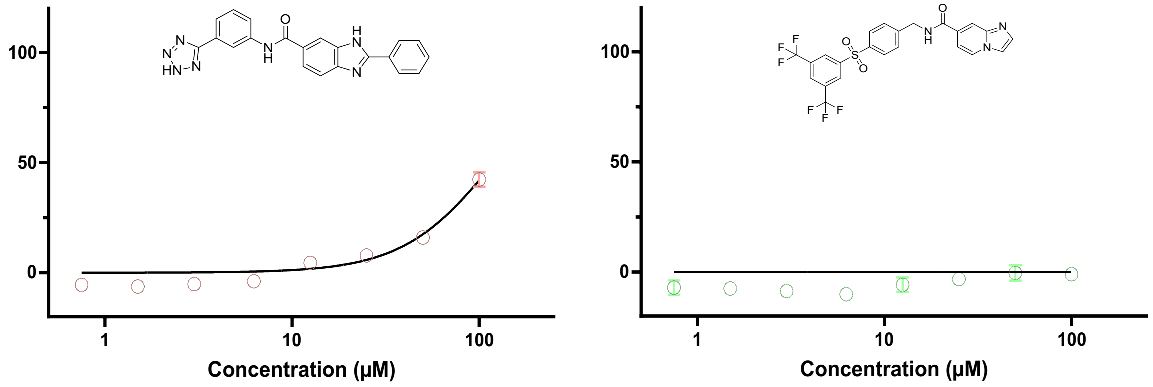


**Figure S3**. The ATP consumption assay for GDI-10928 and GDI-10931 compounds against PheRS target using PF-3845 and DMSO as positive and negative controls, respectively.

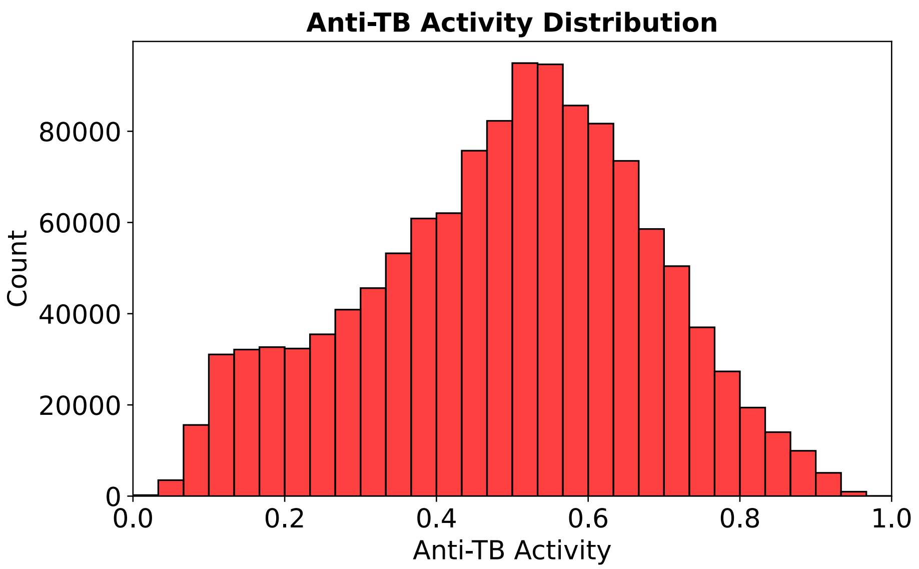


**Figure S4**. Distribution of anti-TB activity probability predicted by Ligandformer model for generated molecules. The predicted score indicates a likelihood that the compound will inhibit M.tb grouth.

s

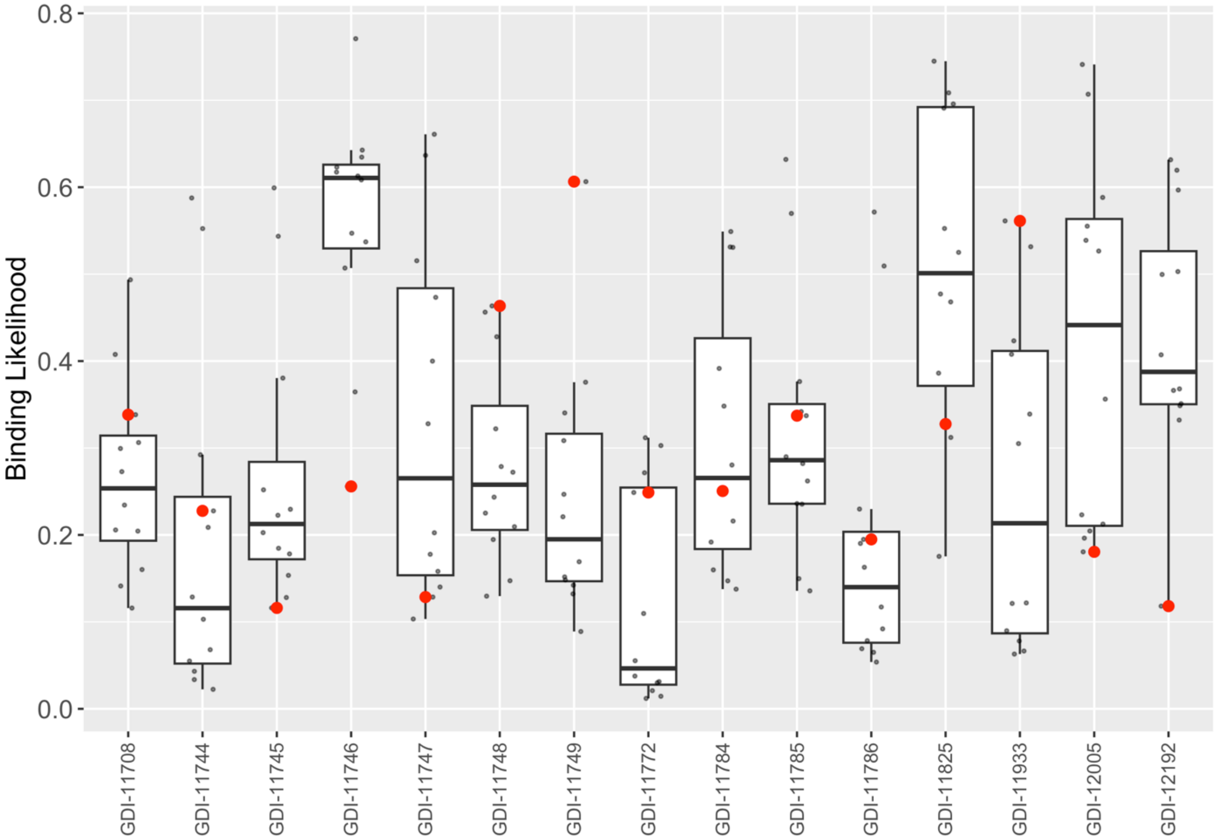


**Figure S5**. The binary binding likelihood for kinase of vero cell. The red points were the compounds with highest docking score at vina.

**Table S1:** The structural and functional annotations of protein pockets identified across Mtb essential targets. Each entry includes: (1) the pocket ID (sequential index), (2) the pocket identifier (e.g., AccA3_5MLK_sitemap), (3) the Protein Data Bank (PDB) or AlphaFold structure identifier, (4) the protein locus tag (Rv number) with its associated gene name in parentheses, (5) Pockets are categorized by their annotation source (e.g., literature, prediction, or homology), (6) the functional description for proteins, (7) the total number of pockets identified per protein, (8) the specific pocket index within the protein, and (9) the three-dimensional coordinates (Å) of the pocket center in the corresponding structure.

**Table S2**. The data source for AI-cellular activity model

| | **Strain** | **Number** |
|---|---|---|
| **GSK** | H37Rv | 177 |
| | H37Rv | 50 |
| | BCG | 599 |
| **GHDDI-Project** | H37Rv (BTTTRI) | 1994 |
| | H37Rv (Broad) | |
| | H37Rv (SCRI) | |
| **GHDDI-HTS** | H37Rv | 1658 |
| **Literature** | CDC1551 | 935 |

MIC values were tested at three different laboratories, BTTTRI: Beijing Tuberculosis and Thoracic Tumor Research Institute; Broad: Broad Institute; SCRI: Seattle Children's Research Institute.

**Table S3.** PAINS counts and percentages in GenVS-TBDB versus known libraries.

| | **Total Count** | **PAINS Count** | **PAINS%** |
|---|---|---|---|
| **FDA** | 2,664 | 68 | 2.6% |
| **Prc&Cli** | 1,129 | 21 | 1.9% |
| **ReFrame** | 11,599 | 244 | 2.1% |
| **GenVS-TBDB** | 1,252,274 | 3298 | 0.26% |

**Table S4**. The TSA assay against MDH target.

| GDI number | Structure | TSA ΔTm (˚C), 1**00/200/400** μM **1**h | Vina score |
|---|---|---|---|
| GDI-10931 | 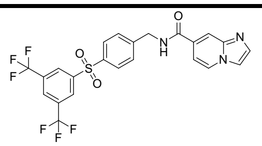 | 0.1/0.2/0.3 | **-10.31** |

The Tm shift is 4.7˚ at 1 mM concentration for positive control NADH.
